# What shapes the continuum of reproductive isolation? Lessons from *Heliconius* butterflies

**DOI:** 10.1101/107011

**Authors:** C. Mérot, C. Salazar, R. M. Merrill, C. Jiggins, M. Joron

**Affiliations:** ISYEB UMR 7205, Muséum National d’Histoire Naturelle, 45 rue Buffon, Paris; IBIS, Université Laval, 1030 Avenue de la Médecine, Québec, Canada; Biology Program, Faculty of Natural Sciences and Mathematics. Universidad del Rosario. Carrera. 24 No 63C-69, Bogota D.C., 111221. Colombia.; Department of Zoology, University of Cambridge, Downing Street, Cambridge, CB2 3EJ, United Kingdom; Smithsonian Tropical Research Institute, MRC 0580-12, Unit 9100 Box 0948, DPO AA 34002-9998; Centre d’Ecologie Fonctionnelle et Evolutive, UMR 5175 CNRS - Université de Montpellier - Université Paul Valéry Montpellier - EPHE, 1919 route de Mende, 34293 Montpellier, France

**Keywords:** Speciation, Hybrid Sterility, Mate choice, Lepidoptera, Magic trait

## Abstract

The process by which species evolve can be illuminated by investigating barriers that limit gene flow between taxa. Recent radiations, such as *Heliconius* butterflies, offer the opportunity to compare isolation between pairs of taxa at different stages of ecological, geographic and phylogenetic divergence. We carry out a comparative analysis of existing and novel data in order to quantify the strength and direction of isolating barriers within a well-studied clade of *Heliconius*. Our results highlight that increased divergence is associated with the accumulation of stronger and more numerous barriers to gene flow. Wing pattern is both under natural selection for Müllerian mimicry and involved in mate choice, and therefore underlies several isolating barriers. However, pairs which share a similar wing pattern, also display strong reproductive isolation mediated by traits other than wing pattern. This suggests that, while wing pattern is a key factor for early stages of divergence, it is not essential at a higher level. Additional factors including habitat isolation, hybrid sterility and chemically-mediated mate choice are associated with complete speciation. Therefore, although most previous work has emphasised the role of wing pattern, our comparative results highlight that speciation is a multidimensional process, whose completion is stabilized by many factors.

## Introduction

Studies of speciation have long contrasted allopatric and sympatric speciation, speciation through sexual versus natural selection, and ecological versus non-ecological speciation. However, these contrasts do not always reflect the diversity of processes involved in divergence and the challenge is to reach an integrated understanding of speciation [1–4]. Species divergence involves multiple different traits and processes that can lead to reproductive isolation [5]. These include adaptation to local environmental conditions, pre-mating isolation as well as post-mating effects that reduce the fitness of hybrids. To untangle the evolutionary processes at play, it is useful to quantify the relative importance of the factors reducing gene flow between diverging populations [6].

Speciation is a continuous process and we can typically only observe the results of divergence at a specific stage, not the process in its entirety. For instance, incompatibilities between extant species may not reveal the ecological and evolutionary forces initially causing phenotypic divergence [7]. Conversely, ecotypes or subspecies at early divergence may shed light on factors favouring early divergence but speciation is not a necessary outcome [8, 9] and the challenge of speciation with gene flow might not be its initiation but its progression and completion [10]. In that context, a useful way to study speciation as a continuous process is to compare multiple pairs of incipient or closely-related species which vary in their extent of divergence, possibly depicting stages along the so-called speciation continuum [9, 11–14].

With a large diversity of recently diverged species and sub-species, the radiation of *Heliconius* butterflies is an excellent system for studying speciation [15]. Within *Heliconius*, the two sister clades, melpomene clade and cydno clade each contains a large number of local representatives across the Neotropics (Fig 1). They provide replicate pairs of sister-taxa distributed along the speciation continuum, notably spanning the “grey zone of speciation” [14], providing an opportunity to assess the factor shaping reproductive isolation along the speciation process.

**Figure 1:**
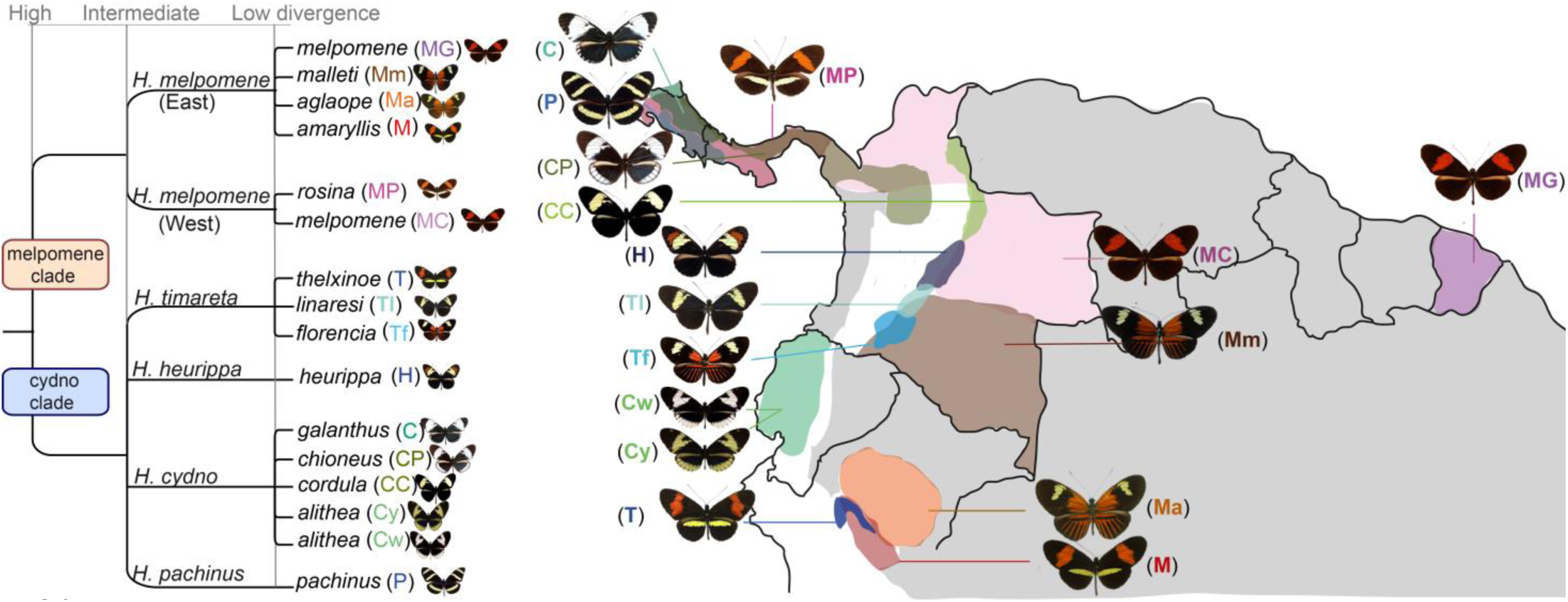
Geographic range and relationships of the taxa included in this study. *H. m. melpomene* and *H. m. malleti* have a wide range through South America but we chose to represent only their range in the country where they were studied. Grey areas represent areas harbouring other subspecies of *H. cydno/timareta* and *H. melpomene* which we did not include in our analyses. Phylogeny is adapted from [16, 17]. Range localisation is adapted from [52].

*H. melpomene* is considered a single taxonomic species but comprises populations with significant genetic differentiation between western and eastern populations on either side of the Andes [16, 17]. The cydno clade includes four taxonomic species, *H. cydno, H. pachinus*, *H. timareta* and *H. heurippa*. Across their range, representatives of the cydno clade are typically broadly sympatric with *H. melpomene* and hybridize at low frequency [18–20], resulting in persistent inter-specific admixture across the genome [21].

Within this clade, the modalities of reproductive isolation have been examined across numerous studies considering taxa at various levels of divergence, falling into three main categories. First, pairs of taxa at low divergence, such as sympatric forms of *H. cydno alithea* (Ecuador) [22, 23] and parapatric subspecies of *H. timareta* (Colombia) [24, 25] or *H. melpomene* (Peru) [26]. Secondly, at intermediate divergence, populations belonging to the same clade (either cydno or melpomene) but characterized by significant genetic clustering. Within the cydno clade, those correspond to separate species, such as *H. cydno galanthus* and *H. pachinus* (Costa Rica) [23, 27] or *H. cydno cordula* and *H. heurippa* (Colombia) [28, 29]. Within the melpomene clade, this includes allopatric subspecies of *H. melpomene* belonging to the eastern and western lineages (Panama/French Guiana)[30]. Thirdly, comparison of highly-divergent pairs, involving a population of *H. melpomene* and a representative of the cydno-group, found in sympatry, parapatry or allopatry (Panama, Colombia, Peru, French Guiana) [28–36].

Emphasis has been given to behavioural pre-mating isolation, found to be strong in most pairs of taxa [23, 26, 30, 34]. However, other components of divergence such as habitat preference [37], hybrid fertility [29, 32], hybrid survival in the wild [36] and hybrid mating success [33] have also received some attention. Here, to provide an extensive comparison across the whole clade, we conduct a joint re-analysis of those published data with new data and quantify the contribution to reproductive isolation of each isolating component.

Most studies focus on pairs of species diverging in wing colour pattern. Wing pattern has been termed a ‘magic trait’ causing speciation, because disruptive selection and assortative mating operate directly on the same trait, wing pattern, thereby coupling two key forms of reproductive isolation [30, 36, 38–40]. First, *Heliconius* wing patterns are warning signals under strong natural selection for Müllerian mimicry [41–44]. Individuals not fitting one of the warning patterns recognised by predators suffer a higher risk of predation and there is evidence for selection against immigrant and hybrid wing patterns [36, 41, 42, 45]. Second, wing patterns are also involved in mate-recognition in *Heliconius*, and males typically preferentially court females displaying their own colour pattern [22, 26, 30, 33]. The loci controlling colour pattern appear to be tightly linked to mate preference loci, which may help maintain the association between signal and preference [23, 35]. Consequently, wing pattern divergence causes reproductive isolation both through hybrid unfitness and assortative mating, and in *Heliconius*, speciation is indeed frequently associated with a colour pattern shift [38, 46, 47].

Cases of mimicry between sibling species were unknown in *Heliconius* until the discovery of new cryptic subspecies of *H. timareta* in sympatry with its co-mimic *H. melpomene* in Colombia, Ecuador and Peru [20, 31, 48, 49]. Less is known about the mechanisms responsible for reproductive isolation between these species pairs with similar wing patterns (co-mimics), but this will be important in understanding the role of mimicry shifts in reproductive isolation. Indeed, wing-pattern similarity may be predicted to favour hybrid matings as well as increase the survival of hybrid adults, and so may weaken both pre-mating and post-mating isolation between taxa.

In this study, we investigate the mechanisms involved in the build-up of reproductive isolation by means of a large-scale, comparative analysis on this clade of *Heliconius* butterflies. We combine new data with data collected from the existing *Heliconius* literature. The numerous studies of *Heliconius* taxon-pairs at various levels of divergence and in various geographic contexts allow us to evaluate the relative importance of different isolating barriers and their appearance at various stages of divergence. We have applied a unified framework for the quantification of isolating barriers that facilitates these comparisons [6]. By contrasting co-mimetic *vs*. non-mimetic pairs of species, we also specifically address the importance of wing-pattern as a ‘magic trait’ for reproductive isolation in *Heliconius*.

## Material and methods

### Species studied

We considered published data from all representatives of the cydno clade, *H. cydno, H.pachinus, H. timareta, H. heurippa* and from the two *H. melpomene* lineages (Fig. 1; see all the references in Table S1 for each pair). New data is provided for the pair of co-mimics *H. t. thelxinoe/H. m. amaryllis, H. t. florencia/H. m. malleti* and three Colombian non-mimetic pairs *H. heurippa, H. c. cordula* and *H. m. melpomene* in supplementary material.

### General framework: quantifying the strength of reproductive isolation (RI)

We quantified the strength of reproductive isolation (RI) for each isolating barrier following [6] inspired by [50, 51]. Briefly, the index *RI* offers a linear quantification of RI associated with the presence of a given barrier relatively to expectations in the absence of all barriers. It allows a direct link to gene flow: *RI* = 1 when isolation prevents gene flow, whereas *RI* = 0 if the probability of gene flow does not differ from expectations without this barrier. Confidence intervals for the index can be drawn from confidence interval on the data.

The strength of RI provided by each pre-mating barrier is estimated with the expression:

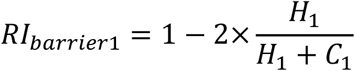

where *H*_1_ is the frequency of heterospecific mating and *C*_1_ the frequency of conspecific mating (when barrier 1 is the only barrier acting).

This is equivalent to:

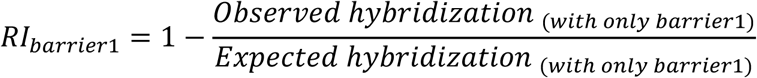

The strength of RI associated with each post-mating barrier is evaluated with the same expression where *H* is the fitness of hybrids and *C* the fitness of pure individuals. RI was calculated separately for both directions of crosses (AxB and BxA; female given first). We summarize hereafter how each barrier was investigated. Detailed methods and calculations are given in supplementary material and Table S1.

### Spatial isolation (at small and large scale)

Since some species ranges are poorly known, the geographic overlap was not evaluated quantitatively. Instead, we provided a qualitative estimate of the level of parapatry, based on the literature [52] in Table S1 and Fig. 1.

Although taxa may overlap in range at a broad geographic scale, this does not necessarily entail equal encounter rates, for instance if relative frequencies differ between microhabitats. For four pairs of species collected in several locations equally distributed along the transition zone (Fig. S1), we use raw collection data (assuming equal collecting efforts on both species) as a proxy for natural encounter rates, and draw an estimate of the expected number of heterospecific vs conspecific matings which we use to calculate reproductive isolation due to spatial distribution, *RI*_spatial._

### Behavioural pre-mating isolating barriers

*Heliconius* males usually patrol the habitat, approach females and perform courtship characterized by sustained hovering and intense wing flapping over the female. Females can accept or reject mating [53]. Most studies have investigated male attraction by visual cues (on models), male preference towards live females, and realised mating. Those three facets of mate choice were analysed separately to dissect their respective contribution to sexual isolation. Realised mating, which reflect the multiple aspects of mate choice by both sexes leading to a mating event, was used for the whole comparison between barriers.

### Model experiments: male attraction on visual cues

In all studies, male preference for different visual cues has been estimated by presenting a male or a group of males with a model made with dead female wings dissected and by recording courtship towards each model.

### Experiments with live females (LF exp): male preference behaviour based on all cues

In all studies, individually-marked males were monitored for courtship during a short time interval when presented with a heterospecific and a conspecific freshly emerged, virgin female.

### Mating experiments: full male-female interaction (M+F)

To investigate mating achievement, most studies have simulated a natural situation, either with a no-choice experiment in which a virgin female (conspecific or hetero-specific) is presented to a one male or a group of ten mature males for 48h, or with a tetrad experiment, where four individuals, one male and one female of each species, were kept until the first mating occurred.

### Post-mating isolating barriers

#### Hatch rate and hybrid sterility

In all studies, mated females of crosses and pure females were kept in individual cages with various fresh shoots of several *Passiflora* species. Eggs were collected on a regular schedule, stored individually in small plastic cups, identified and checked daily for hatching. For each female, fertility was defined as the hatch rate, the percentage of egg hatched over the total number of eggs laid.

#### Hybrid larval fitness

Hybrid survival was recorded only for four pairs (Table S1). In all cases, larvae were raised in individual plastic containers for the first instar. Then, they were gathered by family group in a larger box and fed ad libitum on young shoot of *Passiflora* sp. Survival rate was calculated for each family as the proportion of larvae growing until imago.

#### Hybrid adult fitness

Survival in the wild was tested experimentally only in Panama with paper models of *H. m. rosina/H. c. chioneus* and their F_1_ hybrids [36].

Hybrid ability of mating has been investigated with no-choice experiment or live-virgin female experiment in six pairs (Table S1). For Costa Rican *H. cydno* x *H. pachinus* hybrids, mating success is estimated from experiments with F_1_ wing models.

## Results

### Pre-mating spatial isolation

The highly-divergent species pairs considered here are either sympatric or parapatric but overlap on a large portion of their range. For four of those, local distribution was finely quantified in the area of overlap (Fig. S1). As the probability of encounters between species depends on coexistence within microhabitats, we found that micro-spatial distribution contributes significantly to RI in both mimetic and non-mimetic pairs (*RI_spatial_* =0.7-0.8, Fig. 2, Table 1). For instance, *H. c. chioneus* occupies tall forest habitats where its co-mimic *H. sapho* is abundant, whereas *H. m. rosina* is more frequent in disturbed and edge habitats where *H. erato* is abundant [37, 54]. Similarly, with increasing altitude, *H. t. thelxinoe, H. t. florencia* or *H. heurippa* progressively replace the local *H. melpomene* representative, and are also associated with closed forested habitat.

**Figure 2:**
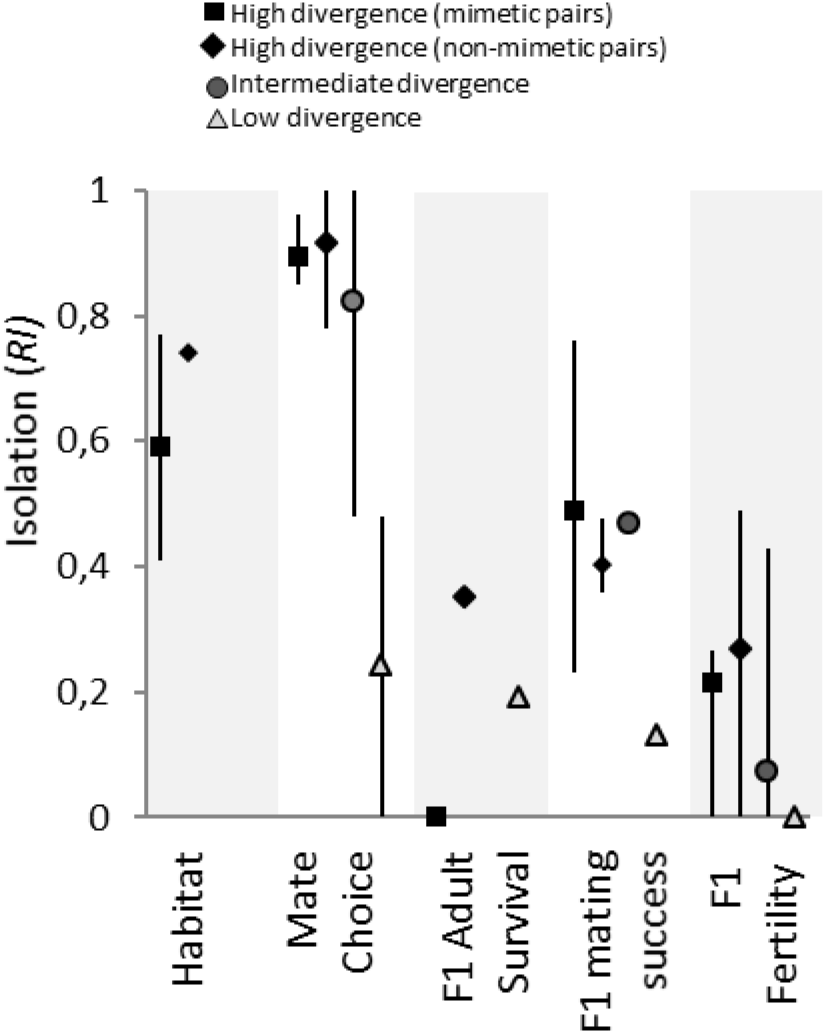
Mean strength of reproductive isolation for each relevant isolating barrier. *RI* associated with each barrier averaged by stage of divergence. The bar range from minimal to maximal values. All detailed values of *RI* are displayed in Table 1

**Table 1:**
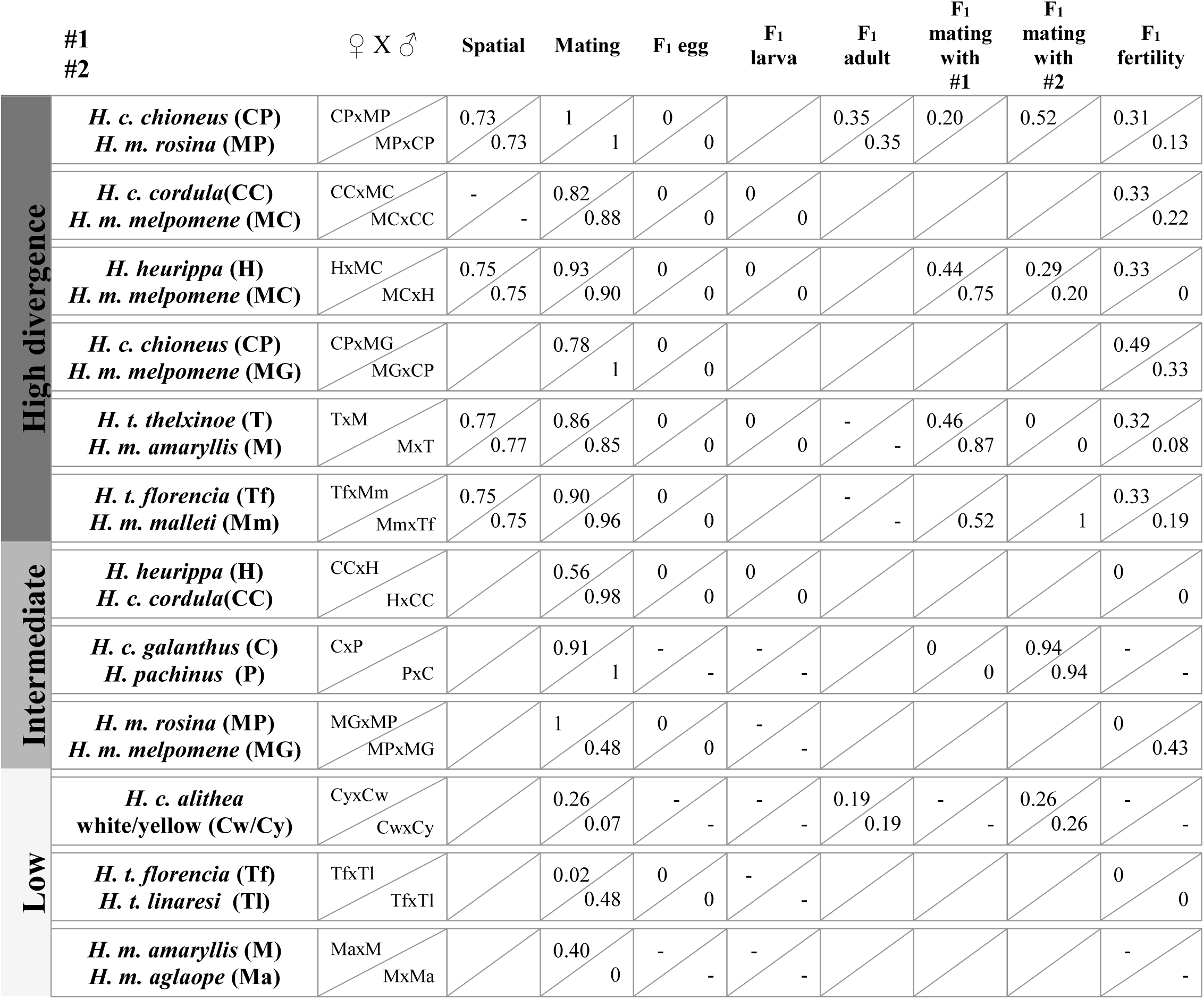
Strength of reproductive isolation associated with each barrier to gene flow. *RI* ranges from 0 (non-significant barrier), to 1 (full isolation). For each pair of species, the two lines correspond to the two possible directions of heterospecific mating with the female given first. Barriers that could not be estimated are not shown. We indicated by a dash barriers that could not be estimated but are likely non-significant.

At intermediate divergence, population pairs are mostly geographic replacements of one another, and do not show any (known) geographic overlap.

At low divergence, *H. cydno alithea* alternative morphs have a patchy distribution, associated with their local co-mimics [41], while pairs of populations within *H. melpomene* or *H. timareta* are geographical races, with narrow hybrid zones where both morphs and hybrids are found.

### Behavioural pre-mating isolating barriers (Fig.3)

**Figure 3:**
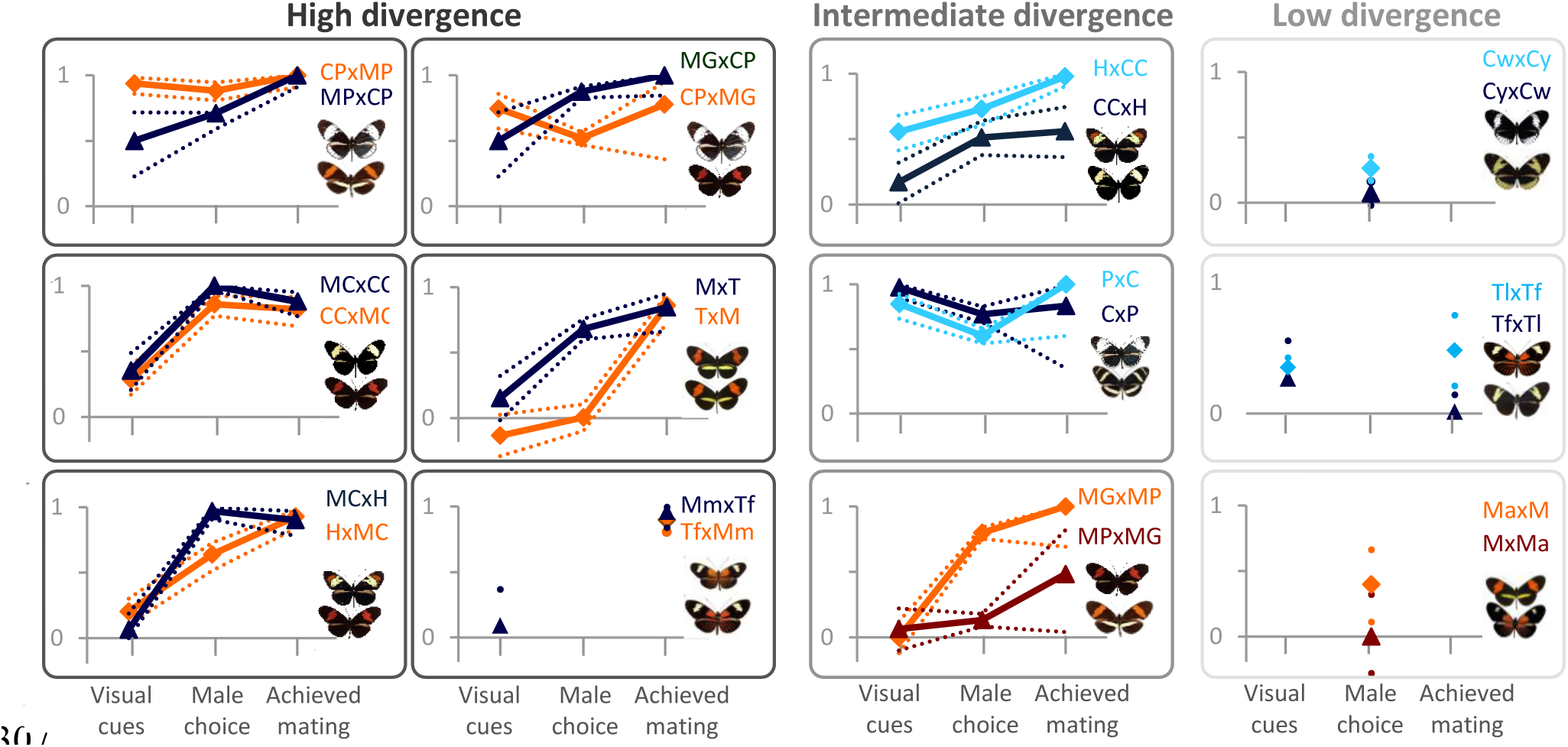
Level of RI associated with each behavioural pre-mating barrier to gene flow. For each pair of species, the two colours correspond to the two possible directions of heterospecific mating with the female given first. Dotted lines are the confidence intervals.

#### Colour pattern model experiments: male preference based on colour

At high divergence, isolation due to male preference based on visual cues is strong for pairs with different colour patterns. It is generally higher in the direction involving *melpomene* males (*RI_colour_* =0.75-0.94, except for the Colombian *H. c. cordula* and *H. m. melpomene* pair reaching only 0.28) than in the other direction (*H. melpomene* X cydno clade males, *RI_colour_* =0.35-0.5). Colour preference is lower between *H. heurippa* and *H. m. melpomene* than between other pairs diverging in colour pattern (*RI_colour_* =0.07/0.2). This might be due to the intermediate pattern of *H. heurippa*, which includes the red band of *H. m. melpomene.* In the co-mimetic pairs (*H. t. thelxinoe* and *H. m. amaryllis; H. t. florencia* and *H. m. malleti*) males do not discriminate between models, as expected given the high visual similarity of the two species.

At intermediate divergence, colour preference remains an isolating factor although its strength varies depending on the pair considered. *RI_colour_* estimates can reach 0.85/0.98 between *H. c. galanthus* and *H. pachinus* but only 0.17/0.56 between *H. heurippa* and *H. c. cordula.* It is zero between the allopatric *H. m. rosina* and *H. m. melpomene*, probably because of the red forewing band shared by the two subspecies.

At low divergence, between *H. t. florencia* and *H. t. linaresi*, some preference is observed, leading to an estimated *RI_colour_* of 0.27/0.35.

#### Experiments with live females: male preference based on all cues

At high divergence, male preference for conspecifics is stronger than in the experiments with models, suggesting that a wider range of proximal cues are available, such as chemical signals or behavioural cues, and influence male courtship decision leading to a higher RI (*RI_male choice_* =0.64-1).

The use of proximal *vs.* long-range visual cues by males seems to depend on the direction of the hetero-specific interaction: *H. melpomene* males appear to choose based on visual cues while cydno-clade males show an accentuated choice with live females, possibly reflecting their response to chemical information (Mérot et al, 2015). *H. melpomene* males indeed respond to wing models with a very strong choice, and appear to show little discrimination when presented with *H. timareta* females showing the same wing pattern, suggesting that additional short-range cues do not play a strong role in *H. melpomene* males courtship decision. By contrast, *H. cydno* or *H. heurippa* males show some discrimination against *H. melpomene* models, but it is weaker than for *H. melpomene* males [30, 35], and choice is generally enhanced by real-females cues. Moreover, in the mimetic Peruvian pair, *H. t. thelxinoe* males strongly prefer conspecific females over heterospecific females using close range chemical cues.

At intermediate and at low divergence, a limited amount of reproductive isolation due to male courtship behaviour is sometimes observed (*RI_male choice_* =0.5-0.78 and 0-0.4, respectively) although the strength of isolation is generally weaker and more asymmetric than at high divergence.

#### Mating experiments: full male-female interaction

At high divergence, the total index of sexual isolation is high for all pairs and in both directions of crosses (*RI_mating_* =0.78-1, Table 1). *RI* estimated using realised mating is higher than when estimated based on visual cues or short range cues from virgin females, suggesting that female response and contact interactions (beyond male courtship) also contribute to pre-mating isolation, especially for the mimetic pairs (preventing *TxM* heterospecific mating for instance).

At intermediate divergence, isolation is generally high, though asymmetric, such as between *H. c. cordula* and *H. heurippa* (*RI_mating_* =0.56/0.98) or between Panamanian and Guianan allopatric populations of *H. melpomene* (*RI_mating_* =0.65/1). *RI* estimated on total mating is again higher than *RI* estimated on experiments with models, suggesting that close-range cues and male-female interactions may also be relevant at intermediate divergence.

By contrast, at low divergence between the parapatric races *H. t. florencia* and *H. t. linaresi*, reproductive isolation is much lower. It is observed only in one direction (TnxTf, *RI_mating_* =0.48) and largely explained by colour pattern preference.

### Post-mating isolating barriers

#### Hatch rate of F_1_ hybrids: cytoplasmic incompatibilities

At high divergence, F_1_ hybrids show no significant reduction of hatch rate (Table S1, S2).

#### Larval fitness of F_1_ broods: host-plant adaptation

Oviposition preferences for different *Passiflora* hosts generally constitute an axis of differentiation between the melpomene and the cydno clade, *H. melpomene* being generally more specialised than its local cydno-clade counterpart [20, 31, 55] with some exception in Colombia where *H. melpomene* has a diverse range of oviposition plants [56].

Hybrid larval survival has only been tested in three pairs at high divergence but shows no significant reduction of survival, leading to a null contribution to reproductive isolation. This suggests neither hybrid viability breakdown related to genetic incompatibilities nor incapacity to metabolize the host-plant are acting in these pairs. For the experiment on hybrids between *H. c. cordula/H. m. melpomene* and *H. heurippa/H. m. melpomene* (Table S5), this result corresponds to expectations since the hybrids were fed on a common host-plant *(P. oesterdii).* However, this may be surprising for the Peruvian pair *H. t. thelxinoe/H. m. amaryllis*, which was fed on the maternal host-plant (Table S2).

Testing survival in experimental conditions with unlimited access to food, fewer parasites and no competition might have underestimated the importance of efficient host-plant use in hybrid growth. We can note for instance, that, in semi-natural conditions, early stage *H. melpomene* larvae from central America had a higher survival rate on *P. menispermifolia* than on other *Passiflora* species [55] while in insectaries, similar growth rates have been achieved for various species of *Passiflora* (Smiley 1978). In Peru, several preliminary attempts of feeding *H. m. amaryllis* larvae and some hybrids (back-crosses towards *H. m. amaryllis*) with *P. edulis* or *P. granadilla* (well-accepted by *H. t. thelxinoe*) led to higher mortality rate.

#### F_1_ adult survival

Adult mortality due to predation was estimated only for the hybrids between *H. c. chioneus x H. m. rosina*. Its contribution to isolation was significant with *RI =* 0.35, but lower than that due to pre-mating barriers.

In the co-mimetic pairs, F_1_ hybrids are visually similar to the parents and predation is not expected to participate in reproductive isolation.

In other cases, F_1_ hybrids may also be similar to one parent (*H. c. galanthus* X *pachinus* hybrids being like *H. c. galanthus* [23], *H. heurippa* X *H. m. melpomene* hybrids being similar to *H. m. melpomene* [29], and heterozygotes at the K locus determining white or yellow morph of the polymorphic taxa *H. cydno alithea* are white (Chamberlain et al, 2009)), which introduces asymmetry in isolation because they are expected to survive better in one habitat. For instance, mark-resight experiments on *H. cydno alithea* (Kapan, 2001) let us estimate predation against white morphs in areas dominated by the yellow mimic, suggesting a mean *RIadult survival* due to predation against F_1_ hybrids around 0.19 (0.38 in areas dominated by yellow, 0 in areas dominated by white).

#### F_1_ mating ability: sexual selection against F_1_ hybrid

At high divergence, in non-mimetic as well as co-mimetic pairs, mate discrimination against F_1_ hybrids appears as an additional isolating barrier although its strength is highly variable and asymmetric, depending on which parental partner is tested (*RI_F_1_success_* =0-0.87, Table 1, Table S3-4, S6-7). At intermediate divergence, mating discrimination against F_1_ hybrids is also observed for *H. c. galanthus x H. pachinus.* Those hybrids are white like *H. c. galanthus* and F_1_ wing models get as much approaches from *H. c. galanthus* males as the pure species, whereas they are discriminated against by *H.pachinus* males, resulting in asymmetric isolation (*RI_F1success_* =0/0.94).

#### Fertility of F_1_ broods: hybrid incompatibilities

At high divergence, the estimated isolating strength of hybrid sterility is intermediate compare to other factors, ranging from 0.33 to 0.49 in one direction and from 0 to 0.33 in the other direction (Table 1, Fig. 2). F_1_ hybrid males are fully fertile except for *H. c. chioneus* (Panama) X *H. m. melpomene* (French Guiana) males which show a slight reduction in fertility [32]. Female F_1_ fertility is more complex. All studies involving crosses between a *H. cydno/heurippa/timareta* mother and a *melpomene* father found complete sterility of female F_1_ (Table S2)[29, 32]. In the other direction of crosses, i.e. a *melpomene* mother x *cydno/timareta/heurippa* father, F_1_ fertility is highly variable. At the extremes, all *H. m. melpomene* X *H. heurippa* females tested were fully fertile [29] whereas *H. m. melpomene* (French Guiana) X *H. c. chioneus* (Panama) females were all sterile [32]. For most other pairs, partial fertility was reported [24, 32] (Table S8) with intriguing non-uniform pattern. For instance, in *H. melpomene* X *H. timareta* hybrids, some hybrid females had a lower fertility, compared with pure females, while others were completely sterile and others completely fertile (Table S2).

At intermediate or low divergence, no significant reduction of fertility was found for the parapatric pairs investigated, neither for *H. heurippa* X *H. c. cordula* [29] nor *H. t. florencia* X *H. t. linaresi* hybrids [24]. The exception is the allopatric pair *H. m. rosina* (Panama)*/H. m. melpomene* (French Guiana) since the F_1_ female (and possibly males) hybrids showed a lower fertility [57], resulting in an additional barrier of 0.43 only for one direction (*MPxMG*).

## Discussion

Quantifying reproductive isolation throughout a speciose clade of *Heliconius* butterflies shows that different levels of genetic divergence correspond to marked quantitative and qualitative differences in reproductive isolation. Higher divergence is associated with both the accumulation of additional barriers and the strengthening of a common set of barriers, although some axes of differentiation are quite labile depending on the ecological context.

The diversity of taxa at different levels of divergence and strengths of RI has been characterised as a ‘speciation continuum’. This does not necessarily imply that these actually represent sequential stages in speciation, nor that any particular example is on an inevitable path towards complete speciation. For example, different stages might be at equilibrium between divergence and gene flow or correspond to qualitatively different pathways to differentiation. Nevertheless, the ‘speciation continuum’ is useful and perhaps analogous to the manner in which those studying the evolution of complex structures, such as the eye or the flagellum, infer past evolutionary trajectories from the comparative study of apparently intermediate structures in extant animals. Such examples provide support for the plausibility of a particular route towards a complex structure, or in the present case a route towards complete speciation, but do not prove that any particular evolutionary route has been taken in nature. Our analysis therefore allows assessment of the roles that different factors might take in shaping divergence, while accepting that the current array of divergence states does not necessarily represent successive stages along a unique path to speciation.

### Is reproductive isolation driven by a single trait or multidimensional factors?

Isolation in the face of gene flow requires that certain factors counter the effects of recombination between alleles that characterise diverging taxa [10, 58–60]. This might include strong disruptive selection on a single (large-effect) trait [61], an association between ecological divergence and reproductive isolation (via a ‘magic’ trait for instance [39]), or the coupling of several isolating barriers [58]. Diverging *Heliconius* taxa showing a shift in colour pattern meet all those criteria, making colour pattern divergence a major initiator and driver of reproductive isolation in this group [38, 62].

Given that colour-pattern differentiation underlies the main isolating barriers (predation, mate choice, habitat partitioning) and that all those barriers operate at low, intermediate and high divergence, one may wonder whether increased isolation results from the “stronger selection” scenario [61], under which barriers associated with colour pattern differences are strengthened along the continuum of divergence. This is the case, for instance, in *Pundamilia* cichlid fish, in which increased isolation is associated with increased divergence on one main axis of differentiation, male coloration in relation with habitat transparency [63]. The alternative hypothesis would be that increased isolation is the product of “multifarious selection” [61], with the addition of independent traits and more isolating barriers at higher divergence [64, 65]. For instance, between colour-pattern races of poison frog, isolation is much higher for a pair which also exhibit size differences associated with habitat specialization [66].

Those predictions can be tested by comparing the strength of the barriers potentially associated with colour pattern divergence along the *Heliconius* continuum. The lower stages of divergence reported in *Heliconius* correspond to wing-pattern races, for which selection causes genetic differentiation only around wing-patterning loci [48] and maintain weak isolation. At this stage, selection on different mimicry associations maintains spatial segregation through predation against migrants [42], and is likely to cause post-mating isolation through predation against non-mimetic hybrids. The third barrier, male preference based on colour, is already acting at low-divergence but its contribution is variable and asymmetric. What is the fate of those barriers at higher divergence? Isolation due to **predation** against hybrids has not been quantified in many pairs of taxa. It does appear stronger for the *H. c. chioneus* x *H. m. rosina* hybrids (high divergence), than for *H. c. alithea* F_1_ (low divergence) for instance. It is worth noting that predation itself is of the same magnitude in both cases, reducing the survival of any deviant form by about 30%. RI due to predation is thus lower in *C. alithea* hybrids because they are similar to one parent (white) while *H. c. chioneus* x *H. m. rosina* hybrids differ from both parents and suffer from predation in all habitats. Therefore, isolation against hybrids depends on dominance and segregation of colour patterns in hybrids, with the hybrid being generally more different at higher level of divergence (except for the mimetic pairs). **Habitat isolation** gets stronger at high divergence. Just like for pairs of taxa at low divergence, fine-scale partitioning between taxa at high divergence may follow the distribution of their co-mimics, as observed for instance between *H. c. chioneus* and *H. m. rosina* across the transition from closed forest to disturbed forest and edge habitat [37]. However, habitat specialization for closed forest habitat is also exhibited by other members of the cydno clade such as *H. timareta* (co-mimic with *H. melpomene*) or *H. heurippa* (which has no co-mimic), suggesting that spatial partitioning at high divergence is not only conditioned by mimicry, but also by other ecological preferences which remain unknown but may involve abiotic conditions, adaptation to altitude or host-plants. The component of **mate choice** clearly attributable to visual cues, deduced from experiments with models, is generally strengthened at high and intermediate divergence, though not consistently between species. In addition, assortative mating is likely to involve a chemical component for most pairs of taxa at high divergence. Again, as hybrids tend to be quite different from parental species at higher divergence, sexual selection against hybrids is also stronger at high divergence. Overall, increased isolation does involve a strengthening of isolating barriers directly linked to colour pattern differences, but higher RI also rests largely on the addition of other isolating dimensions.

To assess the relative importance of colour pattern shift at later stages of speciation, it is also useful to consider species pairs that do not exhibit colour pattern divergence, such as the co-mimics *H. timareta* and *H. melpomene.* Genomic evidence suggests that these species were initially divergent in colour pattern and became co-mimics after secondary introgression of wing pattern alleles from *H. melpomene* into *H. timareta* [67, 68]. Under this scenario, if colour pattern divergence plays an important role in the isolation of species at higher divergence, reproductive isolation is expected to be weakened secondarily by mimicry and gene flow. Such collapse of differentiation has sometimes been observed, notably between pairs of taxa that rely on one main axis of differentiation, habitat-related for instance [69]. Compared with *H. c. chioneus* and *H. m. rosina*, the co-mimics *H. t. thelxinoe* and *H. m. amaryllis* indeed display a ~2% reduction of total estimated RI and a slightly lowered genomic divergence [21]. Both in the Colombian and Peruvian mimetic pairs, natural hybrids are also marginally more frequent (1-3%) [20, 31, 70]. This reduction of RI between co-mimics follows the prediction but shows that lifting the wing-pattern barrier has a rather limited effect on species differentiation because RI relies on multiple other isolating mechanisms (habitat specialisation, assortative mating based on chemical communication [34], partial hybrid sterility and likely host-plant divergence). This implies that reproductive isolation between pairs at high level of divergence is strong enough to allow the secondary loss of certain barriers to gene flow, in this case through introgression of wing-pattern alleles, without compromising genome-wide differentiation. It supports the hypothesis that multiple diverging dimensions add cumulatively to reproductive isolation and favour the completion of speciation in the face of gene flow [61].

### How do isolating mechanisms evolve?

The continuum of reproductive isolation spanned in this study also corresponds to a continuum of time since divergence, raising the questions of how the multiple barriers accumulate through time, which result from selection, which are a by-product of isolation through drift, and what is the relative importance of ecological and non-ecological processes.

Pre-mating sexual isolation stands out as one of the strongest barriers at all levels of divergence and gets stronger along the continuum of divergence. This observation is consistent with the rapid evolution of pre-mating isolation generally reported for speciation with gene flow [2], in fish [9, 71], drosophila [51] or plants [64]. As for darter fish [72], the rapid evolution of strong assortative mating in *Heliconius* appears to be associated with sexual selection, notably for chemosensory traits [73] which, as indicators of mate quality, are common targets of sexual selection [74].

An increase in pre-zygotic isolation between hybridizing populations may also reflect reinforcement, under selection against interspecific mating [75]. In *Drosophila* for instance, the fast evolution of mate choice has been linked to reinforcement processes, with pre-mating isolation being stronger for pairs with geographic overlap [51] and pairs with higher hybridization costs [76]. Here, higher stages of divergence are characterized by a decrease in hybrid fitness, such that stronger pre-mating isolation may reflect stronger selection against hybridization. In addition, the higher geographic overlap seen in pairs at high divergence also provides more opportunities for selection against hybridisation to operate. Evidence for reinforcement comes from higher pre-mating isolation observed in the sympatric *H. c. chioneus/H. m. rosina* than in the allopatric *H. c. chioneus/H. m. melpomene* (French Guiana) as well as an increased mate choice between *H. c. galanthus* and *H. pachinus* in populations close to the contact zone [27].

Under a hypothesis of reinforcement, premating isolation comes as a response to hybrid unfitness, so it may seem paradoxical to observe rather weak or moderate post-mating barriers. It could be that their current contributions do not reflect their past importance or that the accumulation of several weak barriers is sufficient to select for assortative mating. Our analysis may also underestimate the strength of extrinsic post-mating barriers, which are experimentally more difficult to assess. Notably, little is known about the ecology of hybrids, and poor hybrid performance may represent a significant barrier when parental species have markedly different habitat preferences (e.g. altitude between *H. timareta* and *H. melpomene*).

Habitat specialisation associated with fine-scale spatial segregation and host-plant divergence is observed for all pairs at high divergence but for none at low divergence. Interestingly, species at intermediate divergence do not show clear habitat or host-plant differences either, suggesting that habitat specialisation might be one of the key barriers allowing geographic overlap and leading to high divergence. Such a transition from parapatric, ecologically-similar morphs to overlapping habitat-specialized taxa is also reported along the stickleback speciation continuum [9] and perhaps constitutes a tipping point in the evolution of isolation[13].

The last post-mating barrier widely observed at high divergence but generally absent at lower levels of divergence is hybrid female sterility (with the exception of allopatric races of *H. melpomene* [57], at intermediate divergence). This result is quite general in the literature: when speciation occurs with gene flow, post-mating incompatibilities tend to accumulate more slowly than ecological and pre-mating isolation [51, 71, 77], and follow Haldane’s rule by first affecting the heterogametic sex [78]. Generally, the strongest isolation was found between allopatric pairs coming from distant areas (Panama VS French Guiana) whereas in sympatry, F_1_ female sterility can be variable, from fully sterile to fully fertile, suggesting that sterility is variably affected by local gene flow. *Heliconius* female sterility is typically caused by interactions between the Z chromosome and autosomal loci [29, 32, 57]. Among sympatric pairs of taxa at high divergence such as *H. timareta* and *H. melpomene* or *H. cydno* and *H. melpomene*, Z chromosomes are very divergent while autosomes show a strong signal of admixture [21]. Admixture might prevent the accumulation of incompatibilities on autosomes (or may allow its purge following secondary contact), therefore limiting the evolution of female sterility. Such hypothesis would question the stability of this intrinsic barrier, traditionally assumed to be irreversible.

## Conclusion

In summary, we have quantified most of the known components of reproductive isolation across a recent adaptive radiation. Contrasting pairs of hybridizing taxa showing different levels of divergence suggests that speciation involves the strengthening of some isolating barriers but, importantly, seems to require the accumulation of additional barriers. Indeed, the synergistic action of wing pattern shifts and other isolating mechanisms appears to be important for reproductive isolation in *Heliconius*, especially at early stages of divergence. Nevertheless, the case of co-mimetic hybridizing species reveals that certain isolating barriers, and especially wing pattern differences, may in fact be quite labile or partially reversible. This shows that a seemingly key factor in the early stages of differentiation may have its role taken over by other barriers at later stages of divergence. A key promoter of the stability and completion of species divergence thus appears to be the multidimensionality of reproductive isolation.

## Author's contributions

CM and MJ designed the analysis, and the study of the Peruvian taxa. CM performed the meta-data analysis and the new data acquisition on the Peruvian species. CS performed the new data acquisition on the Colombian species. RM and CJ studied the Panamanian species. CM and MJ wrote the manuscript with contributions from all authors. All authors gave final approval for publication.

## Acknowledgments

This clade-comparison has been made possible thanks to all the experiments and investigations led by *Heliconius* researchers in the past years. We are grateful to them for allowing us to re-analyse their data. We thank the Museo de Historia Natural de Lima, the Ministerio de la Agricultura, the SERNANP-BPAM and PEHCBM-ACR for collection and export permits in Peru. We thank Marianne Elias for constructive and helpful comments on this manuscript.

